# S-BEAM: A Semi-Supervised Ensemble Approach to Rank Potential Causal Variants and Their Target Genes in Microglia for Alzheimer’s Disease

**DOI:** 10.1101/2022.11.01.514771

**Authors:** Archita Khaire, Jia Wen, Xiaoyu Yang, Haibo Zhou, Yin Shen, Yun Li

**Affiliations:** Department of Genetics, University of North Carolina, Chapel Hill, NC, USA; Department of Biostatistics, University of North Carolina, Chapel Hill, NC, USA; Department of Computer Science, University of North Carolina, Chapel Hill, NC, USA; Institute for Human Genetics, University of California, San Francisco, CA, USA; Department of Neurology, University of California, San Francisco, CA, USA; Weill Institute for Neurosciences, University of California, San Francisco, San Francisco, CA, USA

**Keywords:** genomics, Alzheimer’s Disease, microglia, semi-supervised learning, ensemble learning

## Abstract

Alzheimer’s disease (AD) is the leading cause of death among individuals over 65. Despite many AD genetic variants detected by large genome-wide association studies (GWAS), a limited number of causal genes have been confirmed. Conventional machine learning techniques integrate functional annotation data and GWAS signals to assign variants functional relevance probabilities. Yet, a large proportion of genetic variation lies in the non-coding genome, where unsupervised and semi-supervised techniques have demonstrated greater advantage. Furthermore, cell-type specific approaches are needed to better understand disease etiology. Studying AD from a microglia-specific lens is more likely to reveal causal variants involved in immune pathways. Therefore, in this study, we developed S-BEAM: a semi-supervised ensemble approach using microglia-specific data to prioritize non-coding variants and their target genes that play roles in immune-related AD mechanisms. We designed a transductive positive-unlabeled and negative-unlabeled learning model that employs a bagging technique to learn from unlabeled variants, generating multiple predicted probabilities of variant risk. Using a combined homogeneous-heterogeneous ensemble framework, we aggregated the predictions. We applied our model to AD variant data, identifying 11 risk variants acting in well-known AD genes, such as *TSPAN14*, *INPP5D*, and *MS4A2*. These results validated our model’s performance and demonstrated a need to study these genes in the context of microglial pathways. We also proposed further experimental study for 37 potential causal variants associated with less-known genes. Our work has utility in predicting AD relevant genes and variants functioning in microglia and can be generalized for application to other complex diseases or cell types.

## Introduction

Alzheimer’s Disease (AD) is the leading cause of death among individuals over 65 years of age across the world (Tahami Monfared et al., 2022). AD is a neurodegenerative brain disorder characterized by the progressive impairment of cognitive and behavioral abilities. It is the prevalent cause for dementia, the general decline in mental function that is significant enough to interfere with daily life. Moreover, in 2020, the cumulative healthcare costs for the treatment of AD was estimated to be $305 billion (Wong, 2020). However, no cure has been discovered for AD and patients must rely on symptomatic treatments that merely delay disease progression. Therefore, a major aim of current research is to characterize the missing genetic risk for AD development and progression. The heritability of Alzheimer’s disease is estimated to be between 60% and 80% (Gatz et al., 2006), indicating a large genetic predisposition for the illness. Despite large, collaborative, genome-wide association studies (GWAS) to identify these missing loci, a limited number of genes are known to significantly affect the risk of developing AD. A large proportion of genetic variation associated with disease can be attributed to the noncoding genome, such as cis- and trans-regulatory variants (Frydas et al., 2022; Liu et al., 2022). Rather than directly altering gene and protein function, these variants often act through epigenetic modifications. Uncovering the relationship between non-coding variants and the development or progression of clinical disease has the potential to encourage research into novel therapeutics (Vitsios et al., 2021).

Accurately predicting the functional effects of non-coding variants is a nontrivial task. GWAS that identify these associated non-coding variants cannot distinguish between the causal variants and their linkage disequilibrium (LD) tags. Furthermore, GWAS remain underpowered in identifying moderate or small effect size non-coding variants involved in mechanisms of complex disease, because of a limited sample size of neuropsychiatric patients (Visscher et al., 2017). Therefore, it is necessary to integrate functional annotation data with GWAS signals to yield insight into regulatory mechanisms caused by non-coding variation (Amlie-Wolf et al., 2019; Sun et al., 2021). Current computational techniques for post-GWAS analyses rely on statistical and machine-learning based scoring frameworks that assign each variant a probability for functional relevance. Specifically, prioritization tools can arrange a list of candidate genes and variants according to their risk of contributing to disease development based on a collection of evidence sources (Zolotareva & Kliene, 2019). As a single source of data can be susceptible to errors and incompleteness, most tools utilize multiple complementary genomic databases for classification. Frameworks that construct a multidimensional and integrative classification probability are capable of capturing multiple facets of variant function simultaneously (Yang et al., 2014). Yet, the predictive ability is heavily dependent on the type and quality of annotation sources. As non-coding variants often have regulatory roles and act through epigenetic mechanisms, data sources on histone modifications, chromatin accessibility and chromatin contacts can provide valuable measures of a variant’s pathogenic significance. Additionally, although many approaches are successful at identifying risk variants that influence disease, they fail to uncover a link to genes. In contrast to solely variant prioritization, gene-variant pair prioritization is significant for variant interpretation and identifying relationships with disease phenotypes (Cano-Gamez & Trynka, 2020). Expression quantitative trait loci (eQTL) data can assess direct association between markers of genetic variation and gene expression levels. Thus, along with the aforementioned data sources, eQTL information proves effective for this purpose. Conventional machine learning approaches utilize a positive-negative (PN) classification technique that assigns a positive label to known variants and a negative label to unknown variants. This can result in contamination of the training data set because of hidden positives, or undiscovered causal variants. Moreover, obtaining negative samples proves to be difficult because biological data sources rarely store information about the absence of disease-variant relationships. Therefore, models that employ PN approaches can suffer from high false-negative prediction rates (Kolosov et al., 2021). Recently, unsupervised and semi-supervised techniques have demonstrated an advantage in the analysis of non-coding regions because they are not biased by the lack of available labeled training examples (Li et al., 2022). Particularly, positive-unlabeled (PU) and negative-unlabeled (NU) learning are semi-supervised methods suited to genomic classification problems, as they can train a model with limited high-quality labeled data. Various semi-supervised methods have been developed, using different techniques to effectively learn from unlabeled examples. Some approaches first identify reliable negative records in the unlabeled dataset and then train an algorithm from positive, unlabeled, and pseudo-negative labels (Manevitz & Yousef, 2001; Yu et al., 2004). Other approaches employ a traditional binary classifier to distinguish positive samples from weighted unlabeled unlabeled examples (Elkan & Noto, 2008). To address the issues posed by noise in the data, Mordelet & Vert (2011) proposed a bagging approach for transductive PU learning. The model uses multiple biased SVM classifiers that are trained on positive examples and a subset of the negative examples. RESVM (Robust Ensemble SVM) built on this approach by resampling the positive examples and using a bootstrapping technique (Claesen et al., 2015). Unlike fully supervised or unsupervised methods, these approaches can jointly take advantage of experimentally labeled samples and thousands or tens of thousands of unlabeled variants (He et al., 2018). However, there still remains room for significant improvement in this field.

The heterogeneity of cells in the brain necessitates cell-type specific approaches to understanding gene expression. Although cell types are known to be interconnected and disease pathology converges on common pathways, various cell types have diverse functional roles. Causal genes and variants may be distinct across the various neuronal and glial populations of the mammalian brain. In the context of AD, the immune system has emerged as a critical component of the disease’s etiology. Particularly, many studies have investigated the involvement of microglia, the primary immune cells of the central nervous system, in the development of AD (Kosoy et al., 2022; Gerrits et al., 2021; Heneka, 2019). Evidence suggests that microglial phagocytosis prevents the aggregation of toxic amyloid-beta (Aβ) plaques, a hallmark of AD development, through the formation of a barrier that envelops them (Jevtic et al., 2017). Studying AD from a microglia-specific lens is likely to reveal causal variants involved in the regulation of immune mechanisms.

Here, we develop S-BEAM: a semi-supervised bagging ensemble model to prioritize variants and their target genes for AD in microglia. Utilizing microglia-specific data, we employed combined positive-unlabeled (PU) and negative-unlabeled (NU) learning to rank non-coding variants and their associated genes that are most likely to function in microglia in the development of the disease, paving the path for future experimental studies and personalized treatments.

## Methods

### Identification of candidate variants using DeepGWAS

We first identified candidate causal variants by integrating results from the latest AD GWAS study with functional annotations using our recently developed DeepGWAS method (Li et al., 2019). Specifically, we started from the GWAS meta-analysis of AD performed by Bellenguez et al. recently, testing the association between 21,101,114 variants and the disease in 111,326 clinically diagnosed or “proxy” AD cases and 677,633 controls. Using standard GWAS association, based solely on genotype and AD phenotype information, they reported 711 genome-wide significant (p-value < 5 x 10^-8^) variants for AD. In this project, we applied DeepGWAS, a neural network based method, to enhance the number of significant variants in the data published by Bellenguez et al.. By using the DeepGWAS output, we were able to test and rank a larger set of AD-associated variants for functional AD relevance in microglia.

DeepGWAS is a 14-layer neural network that takes functional annotation data as input in addition to GWAS association results. The algorithm utilizes 33 features, consisting of pathogenic scores (CADD-PHRED and FATHMM-XF) and brain-related candidate cis-regulatory regions (cCREs), obtained from various external data sources, and the basic GWAS summary statistics (in our case, results directly published by Bellenguez et al.). Taken in summation, these sources provide additional information about a variant’s association potential for a disease. In the output, each variant was assigned a classification probability and variants that passed the prediction probability threshold of 0.5 were assigned to the positive class. These AD-associated variants enhanced by DeepGWAS serve as our pool of candidate functional variants for subsequent testing and ranking.

### Data Preprocessing

To prepare data for input into DeepGWAS, we performed preprocessing on the 21,101,114 variants reported by Bellenguez et al. and relevant functional annotation data. We obtained summary statistics for association with AD of the 21,101,114 variants from the European Bioinformatics Institute GWAS Catalog under accession no. GCST90027158. All data in the catalog is currently mapped to Genome Assembly GRCh38, whereas all functional annotation data was mapped to an older version: GRCh37. We employed the UCSC LiftOver tool to convert the genome coordinates of the variants from the GRCh38 assembly to GRCh37.

We used bedtools (Quinlan & Hall, 2010) to assess overlap between each variant and each of 25 annotation features (detailed below in the “**Feature Vector Construction**” section). Each annotation feature contains information related to a particular functional element. Variants overlapping with a functional element were assigned a value of 1 for that annotation feature and 0 otherwise. Additionally, we obtained CADD-PHRED scores, FATHMM-XF scores, and eQTL data from external databases (Li et al., 2019). Another two features taken into account by DeepGWAS are the LD score (with any variant) and LD score with variants reported by Bellenguez et al. to be associated with AD (i.e., p-value < 5e-8). Lastly, we obtained data on the 3 basic predictors, p-value, minor allele frequency, and odds ratio, from the GWAS summary statistics.

Of the 21,101,114 variants, we dropped 84,809 from our data, primarily attributed to the removal of duplicates and removal of variants for which data was missing in external sources. Therefore, we performed DeepGWAS enhancement on 21,016,305 variants.

### Development of S-BEAM: A Semi-Supervised Bagging Ensemble for Alzheimer’s Variant Ranking in Microglia

We developed a semi-supervised ensemble classification algorithm (**Figure 1**) to rank AD variants that are mostly likely to be functional in microglia, from the list of AD-associated variants outputted by DeepGWAS. Specifically, we focused on the identification of variants that are non-coding and expressed in microglia cell populations. For model training, we assigned 14 variants to be positive controls and 14 variants to be negative controls based on CRISPRi experiments published in Cooper et al. (2022) or generated by us (unpublished). We used Python (version 3.7.15) and R (version 4.2.1) for all calculations and methods done in this section.

**Figure 1:**
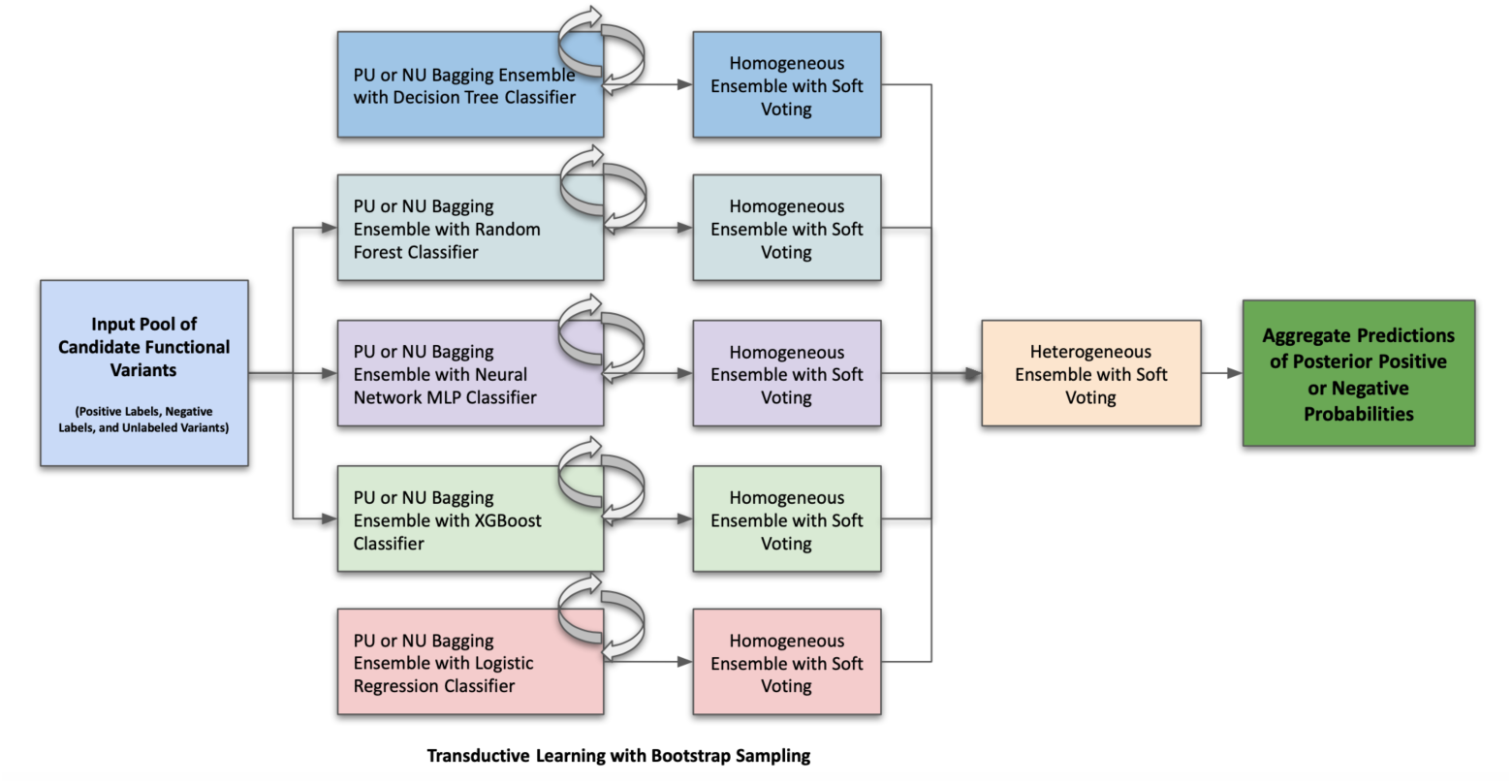
Complete Architecture of S-BEAM. Candidate functional variants identified by DeepGWAS, functional annotation feature vectors, and labeled data are used as input to the model pipeline (leftmost blue box). In the first layer, 5 different homogeneous bagging ensembles are implemented to predict variant posterior probabilities. In the second layer, a heterogeneous voting ensemble takes the probabilities predicted by all 5 classifiers to compute a final probability score using soft voting

### Model Development

We propose a transductive bagging PU and NU learning model (Algorithm 1) with a homogeneous-heterogeneous ensemble framework (Luong et al., 2020) with the goal of computing posterior probabilities of variant functional relevance.

Consider a set of *i* variants with *k* functional annotations for each: *X_i_ = {X_i1_, X_i2_, …, X_ik_}*. We denote a subset *P = {(X_1_, y_1_), (X_2_, y_2_), …, (X_p_, y_p_)}* containing the positively labeled variants, a subset *N = {(X_1_, y_1_), (X_2_, y_2_), …, (X_n_, y_n_)}* containing the negatively labeled variants, and a subset *U = {(X_p+n+1_), (X_p+n+2_), …, (X_p+n+u_)}* containing the unlabeled variants. Additionally, consider a set of *B* different base classifiers. Each base classifier *C* is duplicated *b* times to create a set of identical classifiers: *C_b_ = {C_1_, C_2_, …, C_b_}* and this constitutes a single homogeneous ensemble. Therefore, the heterogeneous module consists of a set of *B × b* classifiers whose predictions are combined for the final output.

Each classifier *C_b_* utilizes both PU and NU binary classification, generating two output probabilities, θ_P_ and θ_N_, for any variant *i* in *U*. To obtain unique training sets for each homogeneous ensemble, we employ a bagging technique (Mordelet & Vert, 2010). A bootstrap sample is created by drawing a subset *U_t_* from *U* by sampling with replacement. We train each classifier *C_b_* to discriminate between *U_t_* and the labeled sets *P* and *N*, then use the classifier to predict θ_P_ and θ_N_ for the out-of-bag set *U_b_* (all *U* not in *U_t_*). This is repeated *k* times, each time drawing a new subset *U_t_* and generating new probabilities θ_P_ and θ_N._ The size *t* of each individual bootstrap sample and *k,* the number of bootstraps, are tunable parameters. θ_P_ and θ_N_ for any variant *i* in *U* are calculated as the sum of θ_P_ and θ_N_ for every bootstrap divided by the number of times *i* is in *U_b_*, denoted by a counter *n*.

#### Algorithm 1. Transudative bagging PU & NU learning.

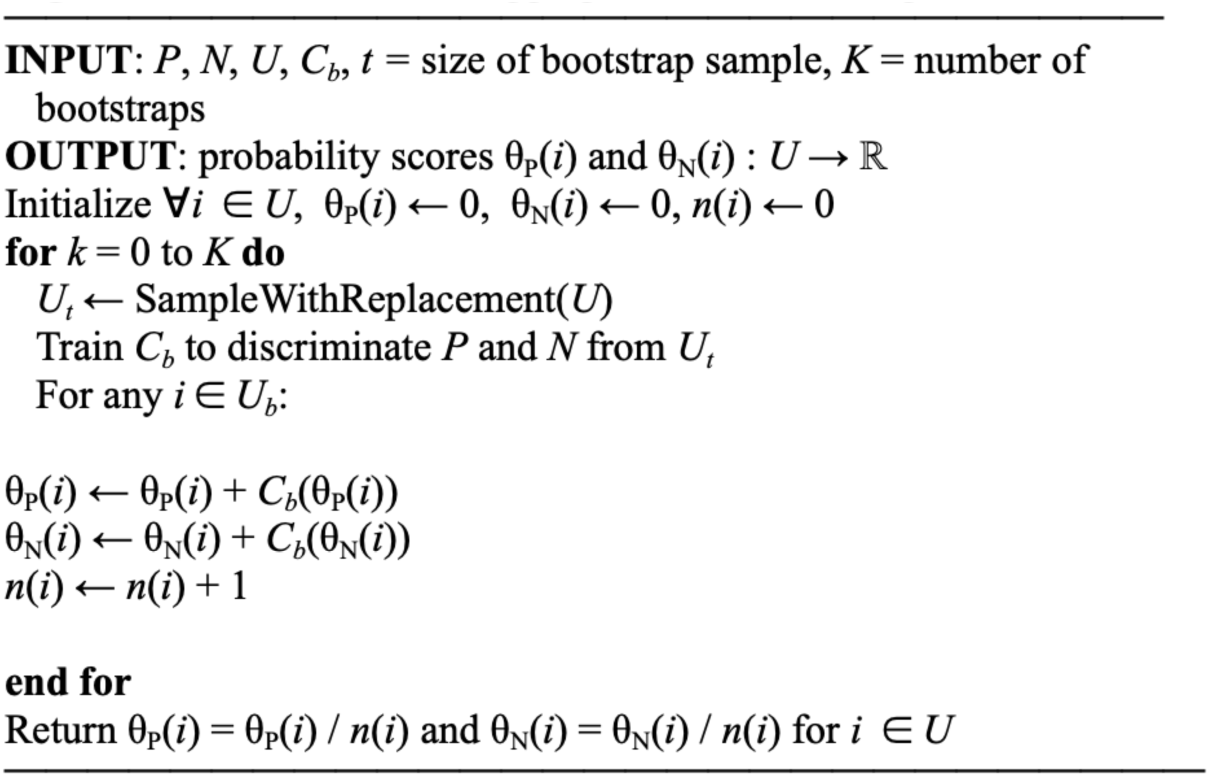

In a single homogeneous ensemble, we generate *j* probabilities θ_P_ and θ_N_ for a specific base classifier *C_b_*, and a soft voting technique is utilized to generate two final probability scores. We then evaluate the agreement between predictions by the set of *b* different classifiers in each homogeneous module. To assess the quality of each base classifier’s performance, we calculated three values: Adjusted Rand Index (ARI), Adjusted Mutual Information (AMI), and Cohen’s Kappa. Classifiers whose predictions were found not to be in agreement with the others were removed from contributing to the probabilities generated by the heterogeneous ensemble. Upon obtaining the list of top-performing base classifiers to constitute the heterogeneous module, soft voting was utilized again to generate two final probability scores, θ_P_ and θ_N_, for any variant *i* in *U*. We set tunable thresholds of θ_P_ > 0.7 and θ_N_ < 0.4 to assign a variant to the positive class.

### Feature Vector Construction

**Table 1** details the 9 functional annotation features that were used in the prediction of a variant’s functional relevance to AD in microglia. We obtained ATAC-seq, H3K27ac ChIPseq, and PLAC-seq data from Nott et al. (2019), eQTL data from Young et al. (2021), and additional ATAC-seq, RNA-seq, and pcHi-C data generated by us (unpublished), following our work in iPSC-derived and primary neurons (Song et al., 2019; Song et al., 2020). To construct feature vector *X_i_* for each variant *i*, we performed preprocessing on this raw data, calculating a functional score for each element.

**Table 1:** Details and Sourcing for the Data Utilized to Generate 9 Annotation Features.

| Feature Name | Details | Description of Functional Significance |
| --- | --- | --- |
| ATAC-seq | Ex vivo; generated from primary microglia | Identifies open chromatin regions |
| ATAC-seq | In vitro; generated from primary microglia | Identifies open chromatin regions |
| H3K27ac ChIPseq | Ex vivo; generated from primary microglia | Identifies regions with acetylation of the histone H3 protein |
| H3K27ac ChIPseq | In vitro; generated from primary microglia | Identifies regions with acetylation of the histone H3 protein |
| PLAC-seq | Generated from primary microglia | Identifies regions with long-range chromatin interactions |
| eQTL | Mapped using RASQUAL | Identifies variants that have significant effects on gene expression levels |
| ATAC-seq |  | Identifies open chromatin regions |
| RNA-seq | Measured in RPKM (Reads Per Kilobase of transcript per million reads Mapped) | Measures RNA transcript quantity |
| RNA-seq | Measured in TPM (Transcript Per Million) | Measures RNA transcript quantity |
| pcHi-C |  | Identifies regions with interactions between promoters and distal regions |

### Model Training on DeepGWAS Output Data

Our unlabeled input set *U* contained 47,702 AD-associated variants outputted by DeepGWAS. We assigned variants that were missing in the feature data to values that were all 0. The set *P* contained 14 confirmed positive variants and the set *N* contained an additional 14 confirmed negative variants. Therefore, we chose a bootstrap size *t* = 14, such that classifiers were trained to discriminate 14 unlabeled variants from *P* and *N*. As we had access to a very limited sample of positive and negative labels (less than 0.05% of the entire dataset), we chose a large bootstrap size of *K* = 4000, such that each variant *i* had a ∼8.04% probability of being in the bootstrap sample.

We utilized *B* = 5 different base classifiers: decision tree, random forest, neural network, XGBoost, and logistic regression. Each homogeneous ensemble consisted of *b* = 3 identical classifiers. Thus, we trained a total of *B × b =* 15 classifiers. These were implemented with the scikit-learn library in Python. We optimized each model separately to select hyperparameters, provided in **Table 2**. For the tree-based methods (decision tree and random forest), we chose a balanced class weight, allowing the algorithm to adjust class weights according to the proportion of class frequencies. When employing a bagging technique, this is effective. We also set the random state to 0, to ensure that model results were reproducible. For the multi-layer perceptron, we utilized two hidden layers with 18 nodes and 5000 iterations, obtained using the sklearn “GridSearchCV” tool. A “relu” activation function and Adam optimizer were also chosen.

**Table 2:**
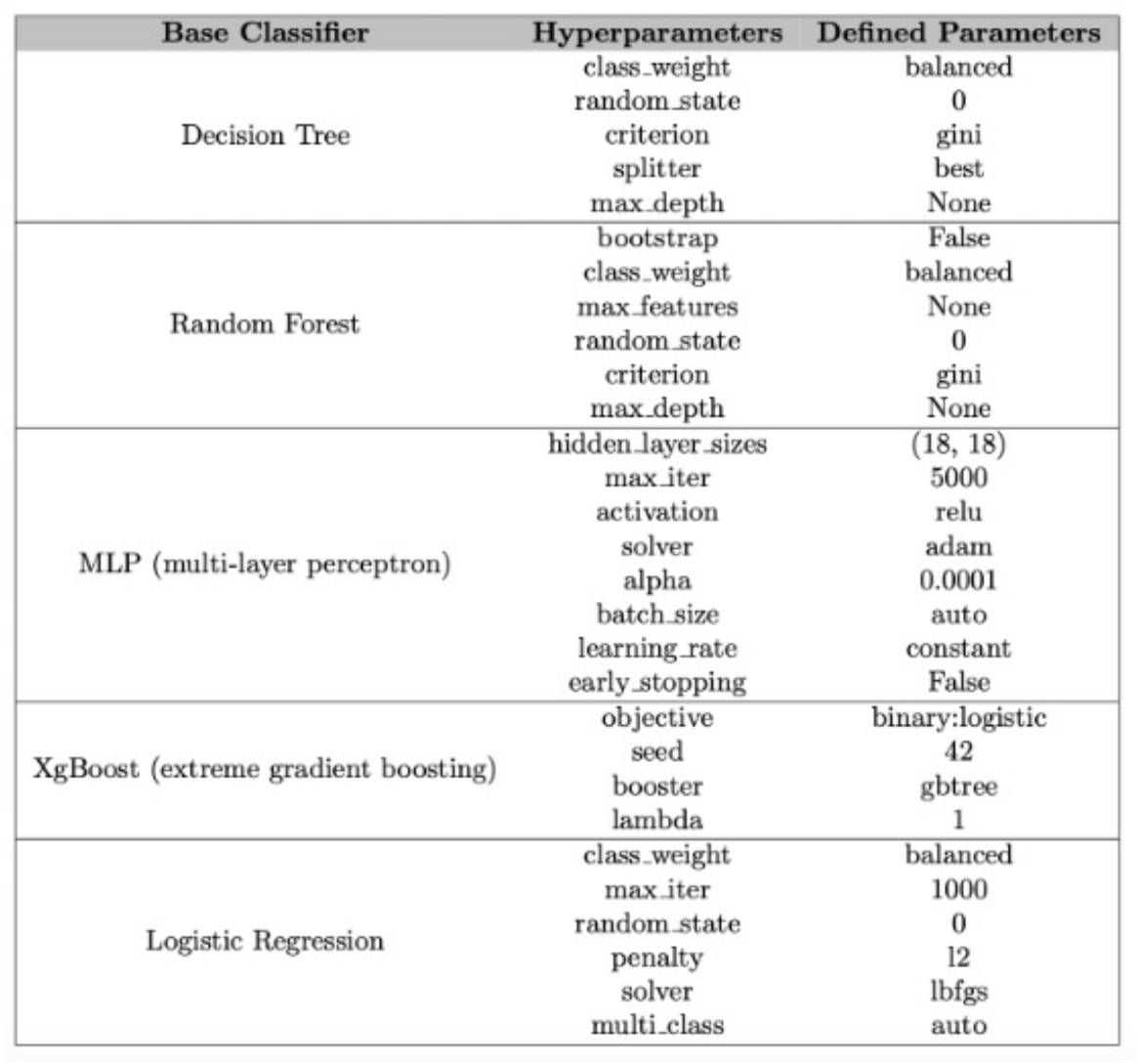
Relevant Hyperparameters Utilized to Train the Five Base Classifiers.

## Results

### Comparison of Results Across Classifiers and Final S-BEAM Output

We set relatively stringent thresholds for the probabilities of belonging to the positive and negative class (θ_P_ > 0.7 and θ_N_ < 0.4). Based on this, we determined that 155 AD-associated variants qualified as potential causal signals with functional relevance (**Figure 2**). To analyze the distribution of variant probability scores, we grouped the output of the PU and NU learning models into 10 bins according to their probability. We found that the distribution in our PU learning model was skewed heavily to the left. 53% of variants were placed in the first bin, where 0.0 ≤ θ_P_ ≤ 0.1, and 16% were placed in the second bin, where 0.1 ≤ θ_P_ ≤ 0.2. The remainder of the variants were relatively uniformly distributed across 0.2 ≤ θ_P_ ≤ 1.0 (**Figure 2b**). In contrast, although the distribution of variants by our NU learning model was also skewed to the left, the difference between individual bins was not as extreme. The first bin, where 0.0 ≤ θ_N_ ≤ 0.1, only contained 20.5% of the variants (**Figure 2c**). This indicates that our PU learning model had stricter thresholds when classifying a variant to be positive than the NU learning model had to classify a variant as negative.

**Figure 2:**
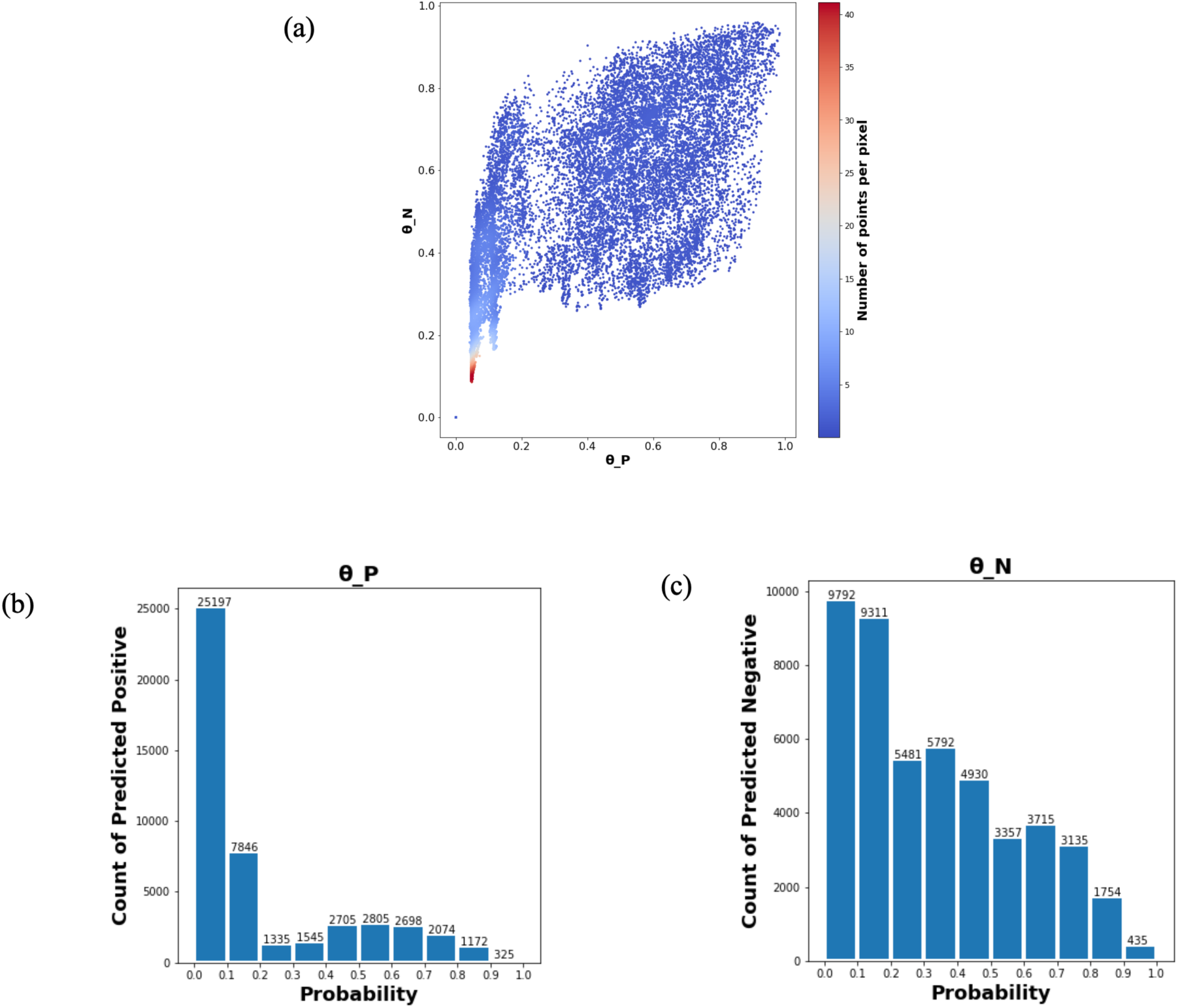
Density Plot of Posterior Positive and Negative Probabilities (θ_P_ and θ_N_) (Top) and Histograms for Predicted Positive and Negative Counts (Bottom) for 47,702 AD-Associated Unlabeled Variants. The density plot depicts a majority of records concentrated between 0.0 and 0.1 for both θ_P_ (X-axis) and θ_N_ (Y-axis), suggesting that the model lacked enough information to classify those variants. The histograms show the distribution of predicted posterior probabilities across 10 bins. We applied thresholds of θ_P_ > 0.7 and θ_N_ < 0.4 to identify the 155 top-performing variants for further analysis. The bins with θ_P_ > 0.7 show the number of variants classified as positive. These are further filtered out with a criteria of θ_N_ < 0.4.

**Figure 3:**
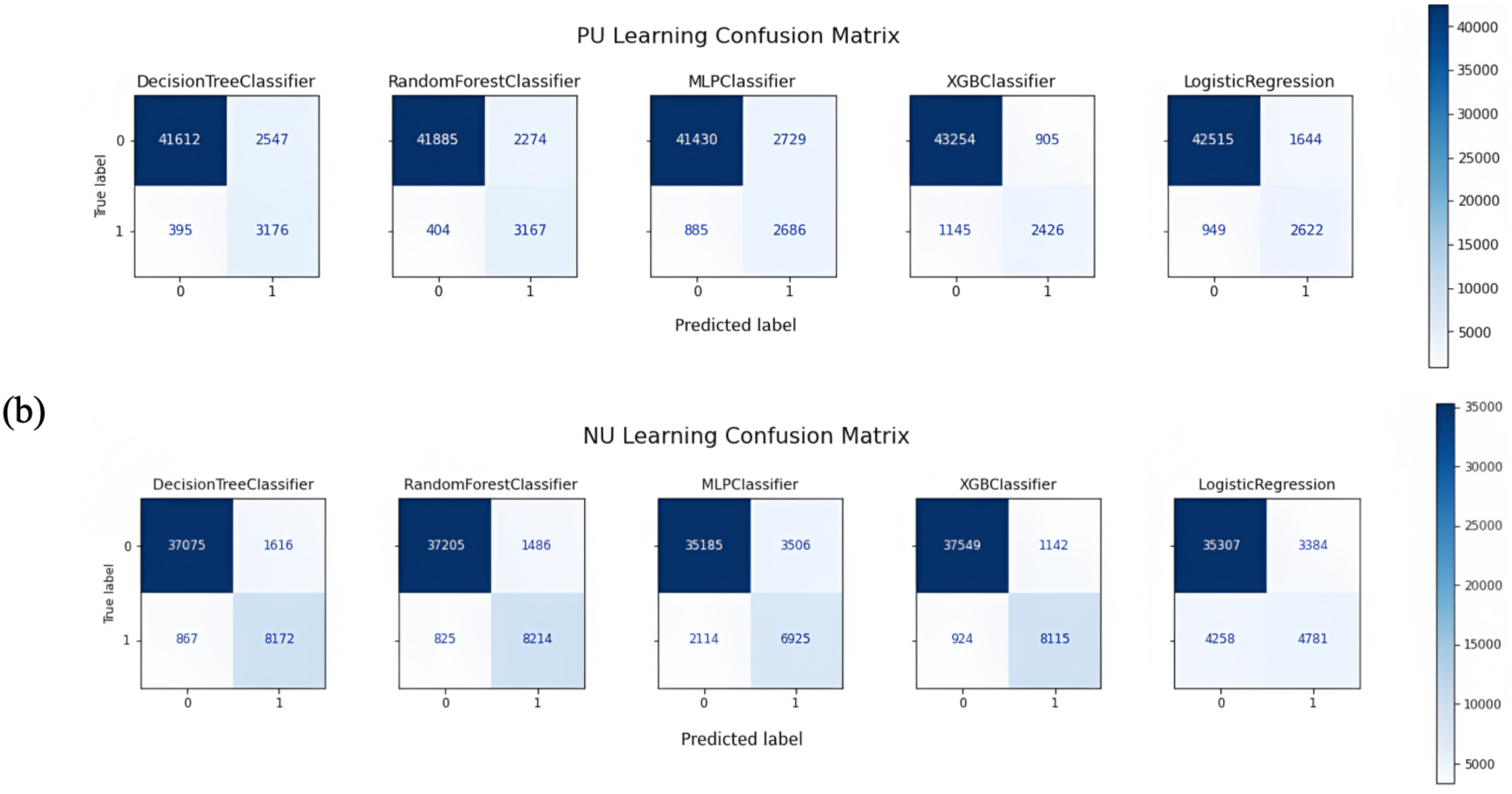
Confusion Matrices for All PU and NU Learning Classifiers. Among the PU learners, decision tree and random forest achieved the highest sensitivities. In the case of NU learning, all tree-based classifiers achieved sensitivities > 90%. Logistic regression had the lowest sensitivity at 55%. The use of a heterogeneous ensemble in our model framework provides the opportunity to drop classifiers that are not in agreement with the others on variant prediction probabilities. We created confusion matrices for each model in PU and NU learning, treating the output of S-BEAM as true positive or true negative labels. For PU learning we used thresholds of θ_P_ > 0.7 and θ_P_ <= 0.7 as the criteria to tag records as true positives and true negatives, respectively. For NU learning, we used slightly less stringent thresholds. Any variant with θ_N_ > 0.6 was treated as a true negative and any variant with θ_N_ <= 0.4 was treated as a true positive. Using these labels, we generated separate confusion matrices for all PU and NU learning classifiers. This enabled an internal evaluation of individual base classifier performance. Particularly, for this study, we were interested in minimizing false negatives and achieving high sensitivity. Among the PU learners, decision tree and random forest achieved the highest sensitivities of 89%, while XGBoost had the lowest sensitivity of 68% (Figure 3a). In case of NU learners, all tree based algorithms (decision tree, random forest, and XGBoost) achieved sensitivities of 90%, whereas logistic regression was the lowest with a sensitivity of 53% (Figure 3b).

We used three different statistical measures, ARI, AMI, and Cohen’s Kappa Score to compare the agreement between our classifiers. For PU learning, all 5 classifiers of heterogeneous ensemble recorded an ARI above 0.5, AMI above 0.3, and Cohen’s Kappa above 0.6, indicating the classifiers were in fair agreement (**Figure 4a**). In the case of NU learning, all classifiers, except logistic regression, recorded an ARI above 0.5, AMI above 0.3, and Cohen’s Kappa above 0.6 (**Figure 4b**). With an ARI of 0.35, AMI of 0.18 and Kappa score of 0.46, the logistic regression classifier in NU learning had slightly lesser scores. Logistic regression uses a log-likelihood objective function to measure accuracy and improve goodness of fit. In NU learning, the imbalanced data may have pushed the objective function to be biased towards the majority class, thus affecting its ability to predict probabilities. For future applications, a potential remedy is to apply penalty weights in order to improve the results of logistic regression. However, we determined that, overall, all of the classifiers were still in relative agreement and performed effectively for our purposes.

**Figure 4:**
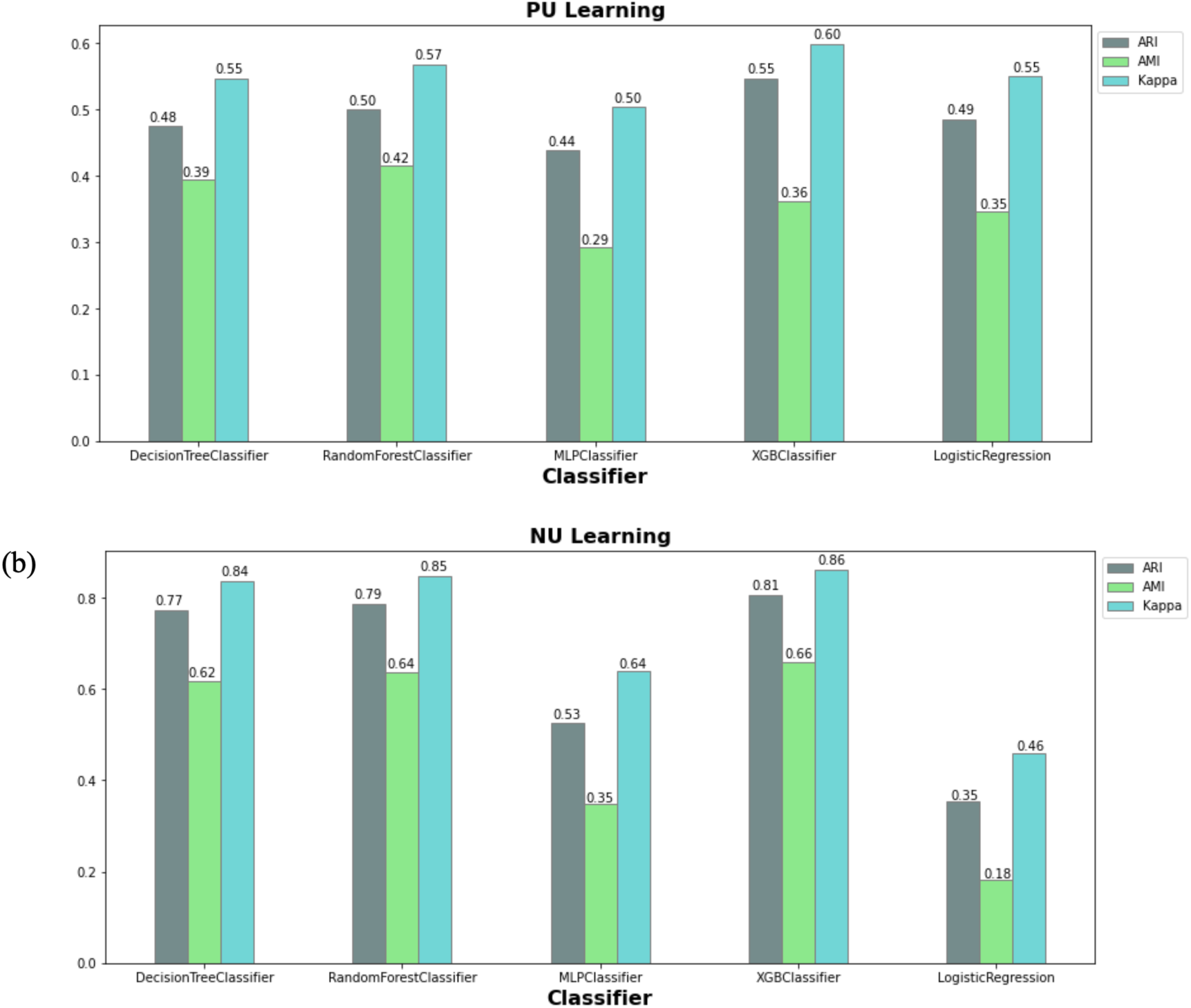
Comparison of 5 Base Classifiers. Three metrics, namely Adjusted Rand Index (ARI), Adjusted Mutual Index (AMI), and Cohen’s Kappa Scores, are compared for the 5 base classifiers in PU Learning (left) and NU Learning (right). All classifiers in the PU learning ensemble achieved an ARI > 0.5, AMI > 0.3, and Cohen’s Kappa > 0.6, indicating fair agreement between all learners. In the case of the NU learning ensemble, all classifiers, except for logistic regression, achieved similar scores. With an ARI of 0.35, AMI of 0.18 and Kappa score of 0.46, logistic regression had less agreement with the other classifiers.

Area Under Precision-Recall Curves (AUPRC) are suited to evaluating the performance of classifiers in the case of imbalanced datasets, which is applicable to disease-gene ranking where there is a limited number of true positives. The baseline value for both the PU and NU auPRCs is ∼0.0003, indicating that the classifiers are performing well with values around 0.5 (**Figure 5**). We can observe that the Random Forest and Decision Tree classifiers suffer from low recall rates at higher decision thresholds for both PU and NU learning. However, they tend to recover to a relatively similar performance to other classifiers at a recall rate of 0.5.

**Figure 5:**
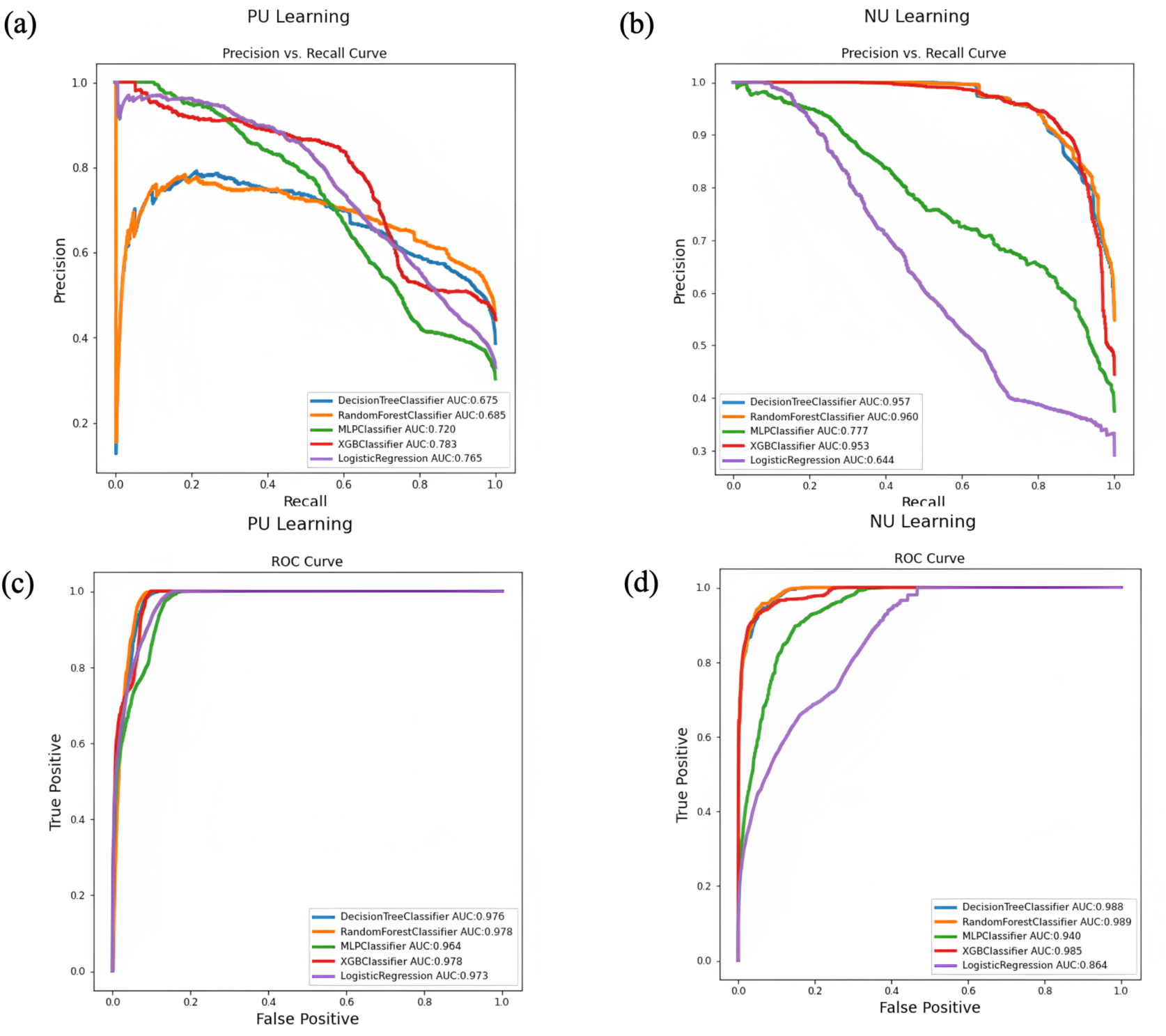
auPRC and auROC for 5 Base Classifiers. Our imbalanced dataset is reflected in the auPRCs. All auPRCs depict high recall and low precision, indicating a limited number of false positives. In disease-variant ranking applications, the objective is to identify some true positives, which our models successfully achieve.

**Figure 6:**
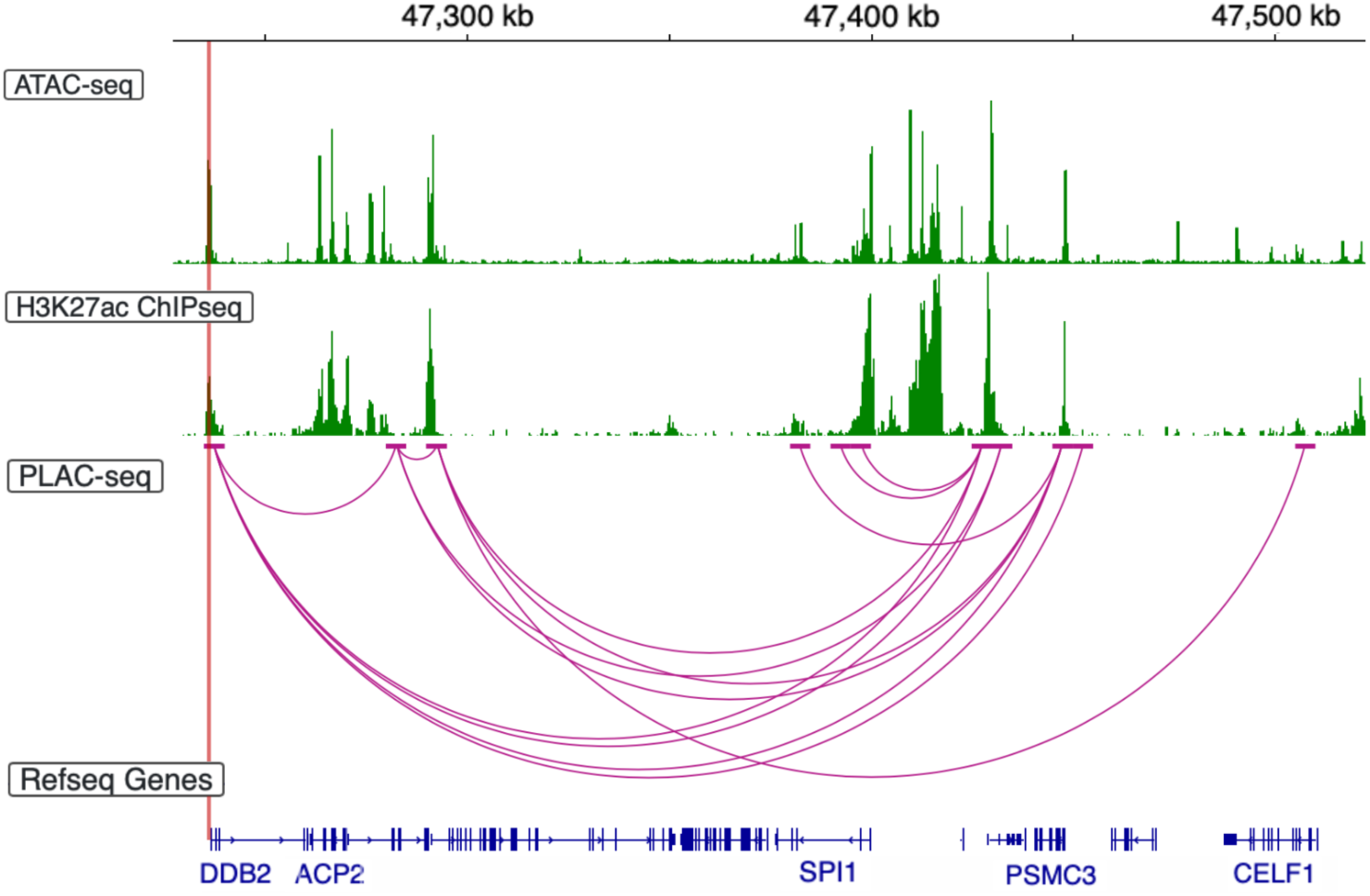
Chromatin Contacts and Open Chromatin Regions near *SPI1* and *CELF1*. As shown by ATAC-seq and H3K27ac ChIPseq peaks and PLAC-seq contacts, interactions can be found between rs11039131, located at chr11:47210487 near *DDB2* and *ACP2*, and known AD risk genes *SPI1* and *CELF1*.

**Figure 7:**
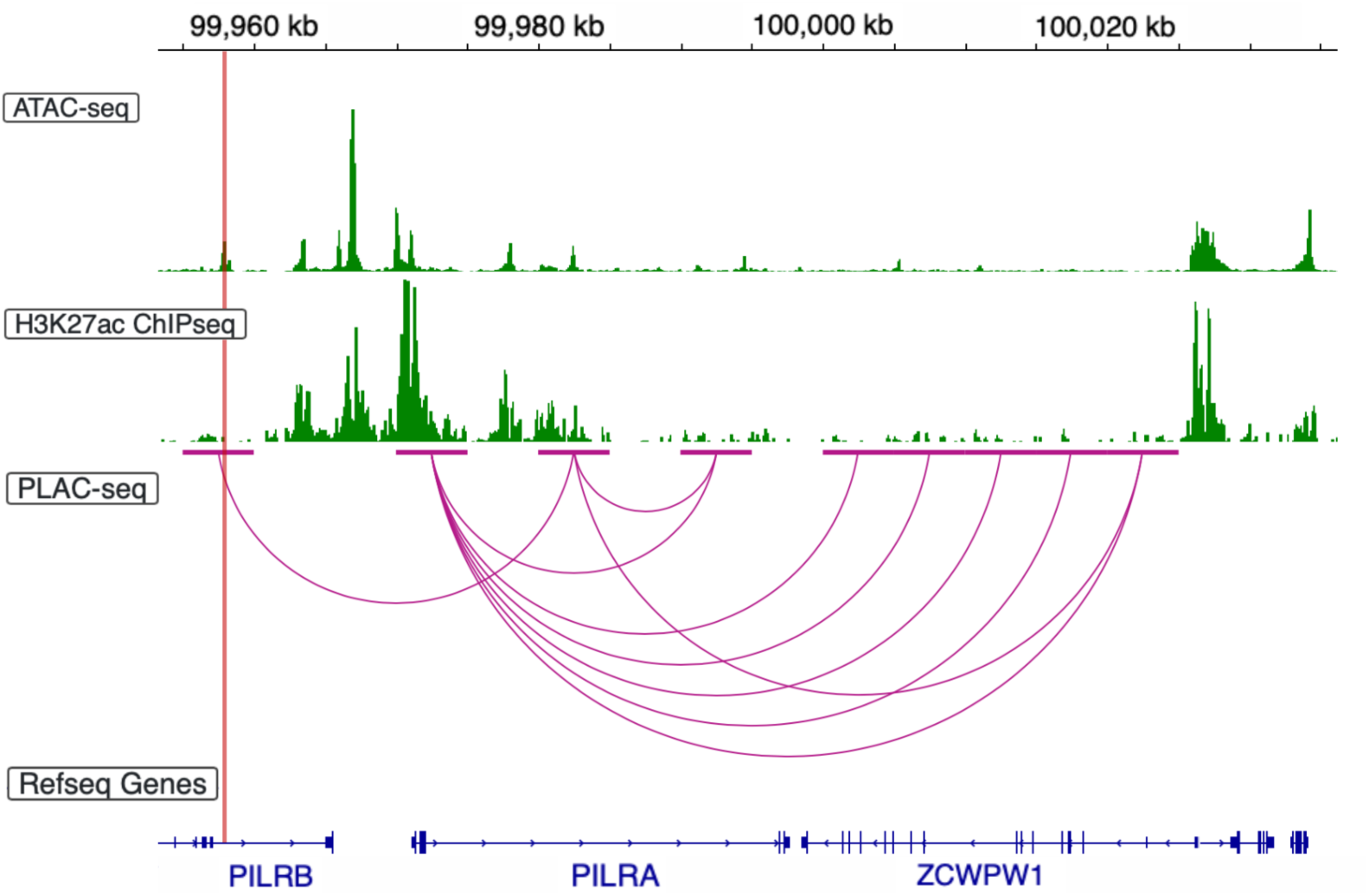
Chromatin Contacts and Open Chromatin Regions near *PILRβ* and the *ZCWPW1* locus. As shown by ATAC-seq and H3K27ac ChIPseq peaks and PLAC-seq contacts, interactions can be found between rs10274982, located at chr7:99984251, and known AD risk loci *ZCWPW1*.

### Top-Performing Variants Evaluated for Potential Causality and Their Target Genes

The lack of existing methods trained on cell-type specific data made it difficult to compare the performance of S-BEAM with external sources. Therefore, we utilized two major criteria to evaluate our top-ranking variants and identify the target genes they were acting on: 1) the variant had chromatin contacts with a relevant gene and 2) previous literature had identified connections between the target gene, microglial mechanisms, and AD. Based on these considerations, we divided the 155 top-performing variants into three groups: 1) variants fulfilling both criteria above and with target genes being well-known AD susceptibility genes, 2) variants fulfilling both criteria above, and with target genes having some research in association to AD and demonstrating relation to microglial activity, and 3) variants and target genes that demonstrated limited or some connection to AD or microglial activity, but fulfilling only one criterion. We proposed that variants which met both criteria were potential causal risk variants and the gene-variant pair warranted further investigation and experimental studies to elucidate their role in AD.

First, we highlight all 11 variants belonging to the first group (**Supplementary Table 1**), those that fulfilled both criteria and whose target genes are well-known AD risk loci. For example, rs10863420 (θ_P_ = 0.85 and θ_N_ = 0.39) and rs12745818 (θ_P_ = 0.73 and θ_N_ = 0.40) demonstrated chromatin contacts with *CR1* and *CR1L*, respectively. rs6737719 (θ_P_ = 0.83 and θ_N_ = 0.36) is located within and has associations with *INPP5D*, a gene selectively expressed in microglia (Tsai et al., 2021). Five variants (rs411920, rs365653, rs2436474, rs12978931, rs554115297) demonstrated chromatin contacts with *NECTIN2*. rs2583476 (θ_P_ = 0.70 and θ_N_ = 0.36) is located within *MS4A2*. rs11755756 (θ_P_ = 0.73 and θ_N_ = 0.36) demonstrated association with *TREML3P*, a gene within the *TREM* locus that also contains the well-established gene *TREM2*. Lastly, rs2343551 (θ_P_ = 0.71 and θ_N_ = 0.38) had contacts with *TSPAN14*. These results validate the performance of our model and also suggest the need for further study of these established genes in the context of immune-related AD pathways.

S-BEAM identified an additional 37 variants (**Supplementary Table 1**) that had been studied previously in relation to AD and demonstrated connections with microglial mechanisms, fulfilling both criteria. We first noticed that *CRHR1*, a receptor that binds to neuropeptides of the corticotropin-releasing family (Futch et al., 2017), was the target gene for 18 of these variants. Although genes for these 37 variants have been less well established for AD, our results suggest prioritizing them for future research in microglia for AD relevance.

Among the 37, we evaluated the roles of 3 variants that demonstrated considerable microglial activity. We first investigated rs11039131 (θ_P_ = 0.82 and θ_N_ = 0.38), which demonstrated significant chromatin contacts with regions containing genes *DDB2* and *ACP2*. *DDB2* is involved with DNA repair and damage response and high rates of DNA damage have been reported in the progression of neurodegenerative diseases like AD (Lutz et al., 2019). These genes are also located within the *CELF1* locus that has previously been identified to contain variants that confer susceptibility for AD (Efthymiou & Goate, 2017). The locus belongs to the *CELF* family of RNA-binding proteins and has roles in the regulation of RNA processing, particularly mRNA stability, splicing, and translation (Ladd et al., 2001). *SPI1* flanks *CELF1* and also contains causal variants associated with AD. It is responsible for encoding a transcription factor, PU.1, that is a master regulator of myeloid cell development and microglial gene expression.

We also analyzed two variants involved in the same AD-associated locus on chromosome 7. rs10274982 (θ_P_ = 0.80 and θ_N_ = 0.39) was located within the gene *AZGP1P1*, which is a pseudogene for *AZGP1*. rs2734895 (θ_P_ = 0.71 and θ_N_ = 0.38) had contacts with *FBXO24* and *PCOLCE-AS1*. Altered levels of the Zinc-alpha-2-glycoprotein, involved in lipid metabolism and encoded by *AZGP1*, were found in AD patient serums (Shen et al., 2017). *FBXO24* encodes a subunit of the F-box proteins that control proteasomal degradation (Omolaoye et al., 2022) and have been linked to the dysfunction of the ubiquitin-protease system, an accepted cause for AD (Watanabe et al., 2013). *PCOLCE-AS1* is likely related to proteases involved in procollagen processing (Podvin et al., 2021), and interestingly, collagen VI was found to counteract Aβ-induced neurotoxicity (Dubal et al., 2008). Furthermore, all four of these genes are located within the *ZCWPW1* locus (Efthymiou & Goate, 2017), a region that is implicated to contain major functional AD genes (Gao et al., 2016).

Specifically, *PILRβ*, a gene neighboring *ZCWPW1*, encodes a paired immunoglobulin-like receptor that is significant in regulating aberrant inflammatory responses (Tato et al., 2012). *PILRβ* is posited to serve as a *DAP12* receptor on microglia, similar to *TREM2* (Rathore et al., 2018). *DAP12* has been identified as a key network hub gene and forms a connection between *TREM2* and *APOE* transcription within microglia. Moreover, absence of *DAP12* was found to decrease microglial recruitment around Aβ deposits. Differential *DAP12* expression also showed evidence of increasing tau phosphorylation (Haure-Mirande et al., 2022). Therefore, our results further implicate the crucial role of genes in the *ZCWPW1* locus for AD pathology and suggest their involvement in microglia and immune-related pathways. We propose that rs10274982 and rs2734895 may be causal AD variants that function through genes *AZGP1, FBXO24,* and *PCOLCE-AS1*, and have a relationship with the *PILRβ-DAP12* pathway.

### Functional Annotation Feature Importance

We analyzed the importance of various features to assess which functional annotation elements had the largest contributions to the predictions made by our model. This was done using the ELI5 package in Python. The feature containing our unpublished ATAC-seq data, obtained from iPSC derived microglia, had the most influence into calculating variant posterior probability (**Figure 8**). In contrast, the features generated from Nott et al. (2019), e.g., ATACseq-exvivo-fetal-MG and ATACseq-invitro-fetal-MG, obtained from fetal microglia, had limited importance (**Figure 8**).

**Figure 8:**
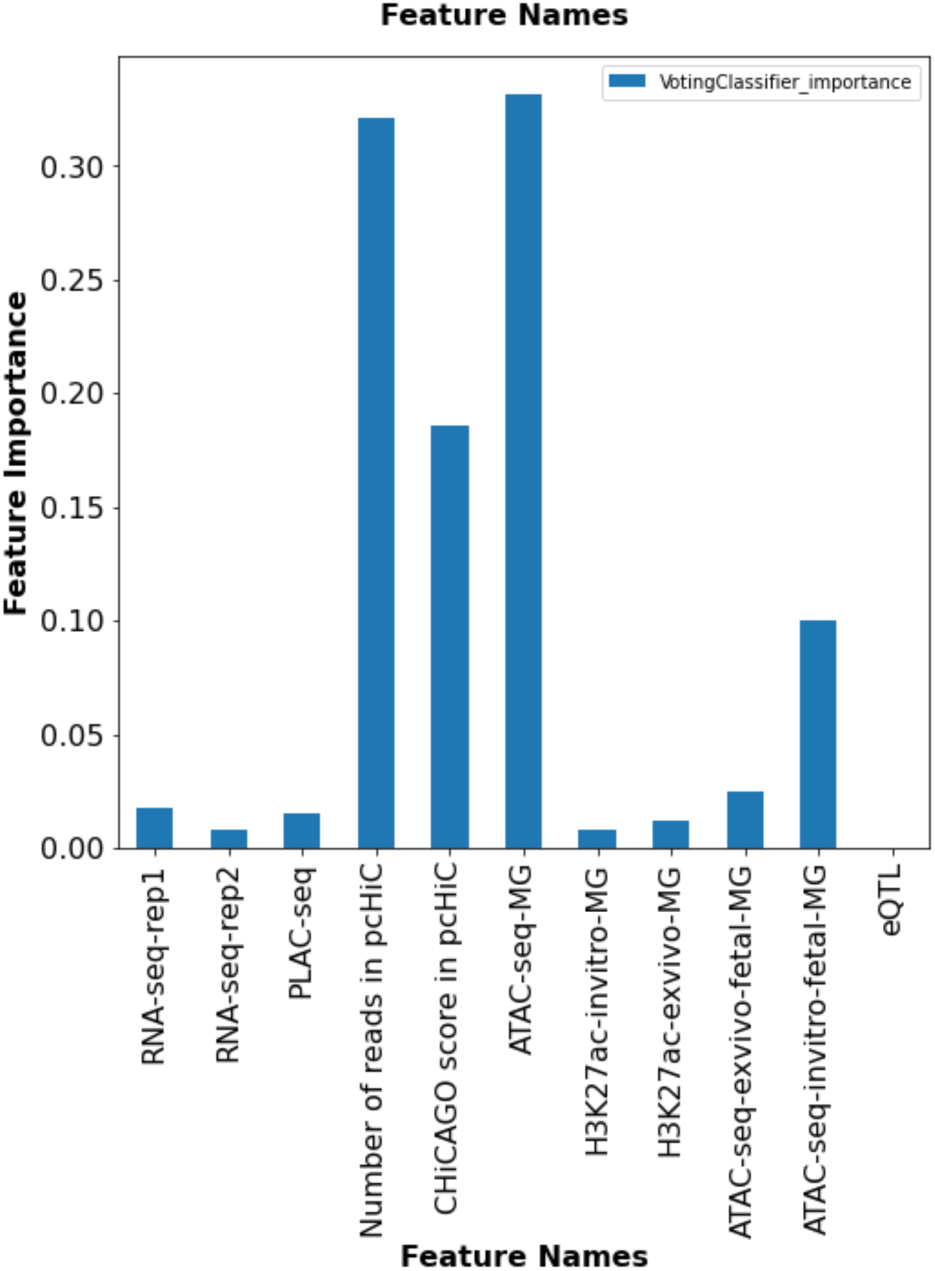
Importance of Features Utilized to Generate Predictions in S-BEAM. Four features are prominent in influencing the decisions made by our model. ATAC-seq data (ATAC-seq-MG), containing information on open chromatin regions, had the greatest influence. This was followed by two pc-HiC features (Number of reads in pcHiC and CHiCAGO score in pcHiC), which capture chromatin contact information.

Given the scarcity of publicly available and preprocessed microglia-specific functional annotation data, we had limited choice of features for our model. Three of our nine features utilize ATAC-seq data, providing insight into whether variants reside in open chromatin regions. Regulatory elements must have a high accessibility to enable binding of transcription factors that activate or silence target genes. Compared to other techniques, ATAC-seq studies are easier to perform across different cell types, making them the most widely used method to quantify chromatin accessibility (Gontarz et al., 2020). RNA-sequencing is an approach to transcriptome profiling that enables interpretation of the functional elements of the genome (Wang et al., 2009), which is a distinct lens compared to the rest of our features.

## Discussion

In this study, we present S-BEAM: a novel bagging PU and NU learning ensemble framework to rank potential causal AD variants by utilizing cell-type specific functional annotation data. We focus on the identification of non-coding variants with regulatory roles that demonstrate associations with microglial genes and immune mechanisms. Our initial set of candidate functional variants was first identified by implementing DeepGWAS, a neural network to enhance GWAS results, on data published by Bellenguez et al. We then trained S-BEAM to further rank these variants according to their probability of pathogenicity, utilizing a combination of ATAC-seq, CHIP-seq, PLAC-seq, RNA-seq, eQTL, and pcHi-C data. Below we discuss several advantages of our method and its application to disease-variant rankings.

Our method utilizes an extension of the PU bagging approach proposed by Mordelet & Vort (2010) that trains a classifier to discriminate positive examples from random subsamples of the unlabeled set. We train a classifier to discriminate both positive and negative examples from random subsamples of the unlabeled set. By treating the existence of positive, negative, and unlabeled examples as two different binary classification problems, we identify with high confidence the probability of an unlabeled variant belonging to the positive class or belonging to the negative class. Unlike most models that employ solely PU learning, utilizing true negative labels serves to reduce the probability of false positives, which can be particularly damaging in disease-variant ranking applications.

Semi-supervised learning and ensemble learning are two prominent paradigms that both attempt to achieve strong generalization by exploiting unlabeled data and employing multiple learners, respectively. Zhou (2009) described the benefits of the combination of both fields, particularly, they find that classifier combination can mitigate the instability caused by training with unlabeled data (Vries & Theirens, 2021). Notably, combined homogeneous-heterogeneous ensembles have demonstrated a better performance than the benchmark of solely one or the other (Sabzevari et al., 2022). In order to overcome the stochastic nature of individual classifiers, we incorporate homogeneous ensembles into our framework. We then pool these different base classifiers to form an overall semi-supervised heterogeneous ensemble and generate diverse predictions.

Our results suggest a potential link between rs11039131 and *SPI1’s* and *CELF1*’s regulation of microglial phagocytosis and anti-inflammatory mechanisms and propose that rs11039131 may serve as an AD risk variant. Research has found that *CELF1* is involved in the regulation of alternative splicing for exon 3 of the *TREM2* locus (Yanaizu et al., 2020), which serves to make a triggering receptor protein that plays crucial roles in AD pathophysiology. *TREM2* regulates microglial migration toward and phagocytosis of amyloid-beta plaques and studies in rodent models have shown that increased *TREM2* expression has reduced amyloid pathology (Zhao et al., 2022). This evidence is further supported given that *CELF1* expression is enriched within microglia. Moreover, the *TREM2*-*APOE* pathway regulates a phenotypic switch from homeostatic to neurodegenerative microglia (Krasseman et al., 2017). Targeting this pathway has potential as a therapeutic treatment to restore homeostatic microglia status.

Altered expression levels of *SPI1* are correlated with phagocytic activity of microglial cells, such that lower expression may reduce AD risk (Huang et al., 2017). PU.1 knock-down renders microglia more susceptible to apoptotic cell death and decreases microglial phagocytosis (Pimenova et al., 2021). It also promoted the anti-inflammatory state of microglia, a protective mechanism in AD. PU.1 has also been noted to bind to *cis*-regulatory elements of several AD-associated genes. Particularly, expression of *SPI1* and PU.1 regulated genes, including *TREM2* and *APOE*, was determined to be elevated in post mortem pathologically confirmed AD tissue (Rustenhoven et al., 2018). Combined, these results encourage further work into investigating PU.1 inhibition as another target for AD treatment. Our results support emerging findings of associations between AD and a network of microglia expressed genes, notably including the well-studied risk factors *TREM2* and *APOE*.

There are several limitations of our study. The power was limited by the lack of experimental data surrounding variants, particularly in a cell-type specific context for AD. Our results reaffirm the need for additional research that generates enough samples to build reliable and useful predictive models. Recent efforts by the IGVF research consortium have made substantial progress in this area, as they work to build a catalog on the impact of genomic variants on function. Despite this, we were able to identify a handful of AD variants whose functional significance was supported in literature, verifying the validity of our approach.

There are further benefits to be found in employing deep learning algorithms as the base classifiers to compose homogeneous ensembles in our method. These techniques have been shown to optimize results because of their ability to capture complex patterns in large datasets, providing a deeper understanding of variant function (Nicholls et al., 2020). However, deep learning also tends to perform poorly when limited data is available, therefore we made the choice to use more traditional algorithms given the data we had access to. In terms of evidence sources, incorporation of sequence-related features may improve the predictive performance of our models. Functional annotation data fails to capture aspects of variant function that can be conveyed through study of DNA, protein, and amino acid sequence conservation (Frousios et al., 2013).

To our knowledge, we are the first to implement a semi-supervised ensemble approach to ranking potential causal disease variants. We hope that S-BEAM has utility in predicting associations between microglia-expressed genes and variants and AD immune mechanisms. We also imagine that our framework can be generalized to identify the impact of variants and genes on other complex diseases, providing basework for future experimental studies and curative therapies.

## Code Availability

The source code to perform data processing and run S-BEAM are available at the following GitHub repositories: https://github.com/akhaire21/Semi-Supervised-Ensemble-Model/blob/main/Microglia_AD_Research_Data_Preprocessing_FinalVersion.ipynb and https://github.com/akhaire21/Semi-Supervised-Ensemble-Model/blob/main/Microglia_AD_Research_PUNULearningModel_FinalVersion.ipynb.

## Biographical Notes

Archita Khaire is a highschool student at the North Carolina School of Science and Mathematics. Jia Wen is a postdoctoral researcher in the Department of Genetics at the University of North Carolina at Chapel Hill.

Xiaoyu Yang is a postdoctoral fellow in the Institute for Human Genetics at the University of California at San Francisco.

Haibo Zhou is a professor in the Department of Biostatistics at the University of North Carolina at Chapel Hill

Yin Shen is professor in the Institute for Human Genetics, Department of Neurology, and Weill Institute for Neurosciences at the University of California at San Francisco.

Yun Li is a professor in the Departments of Genetics, Biostatistics, and Computer Science at the University of North Carolina at Chapel Hill.

## Acknowledgements

We thank the Li lab members for providing advice on data preprocessing, method selection, and feedback on the manuscript.

## Key Points

● Causal mechanisms underlying Alzheimer’s disease (AD) remain poorly characterized functionally, particularly in a cell-type specific manner, because the vast majority of AD associated genetic variants are non-coding.

● We and others have recently generated multi-omics data in microglia for comprehensive prioritization of AD-relevant genes that exert their effects in microglia.

● The small number of known functional variants makes standard supervised machine learning methods based on positive-negative classifications inappropriate because these methods assume unlabeled variants are negative. In contrast, semi-supervised methods that leverage positive-unlabeled (PU) and negative-unlabeled (NU) learning are more suitable.

● We present S-BEAM: a semi-supervised ensemble approach using microglia-specific data to prioritize non-coding variants and their effector genes that play roles in immune-related AD mechanisms.

● S-BEAM prioritized 11 variants in well-known AD genes, such as *TSPAN14*, *INPP5D*, and *MS4A2*. In addition, we further prioritized 37 potential causal variants associated with less-known genes.

---

**Supplementary Figure 1:**
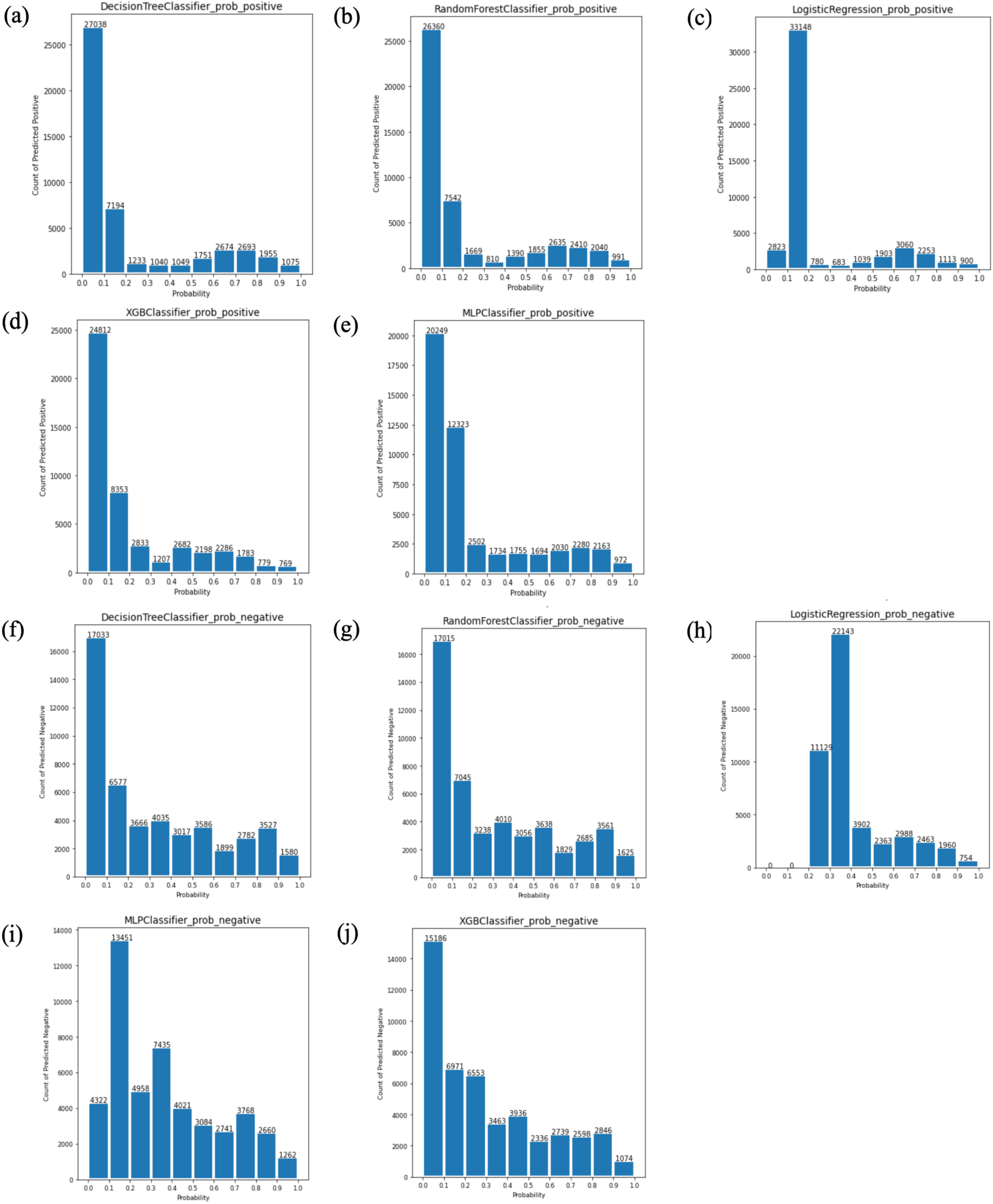
Bin Charts for Individual PU and NU Classifiers. These histograms show the distribution of predicted posterior probabilities across 10 bins for each of the 5 base classifiers implemented in our model.

**Supplementary Table 1:**
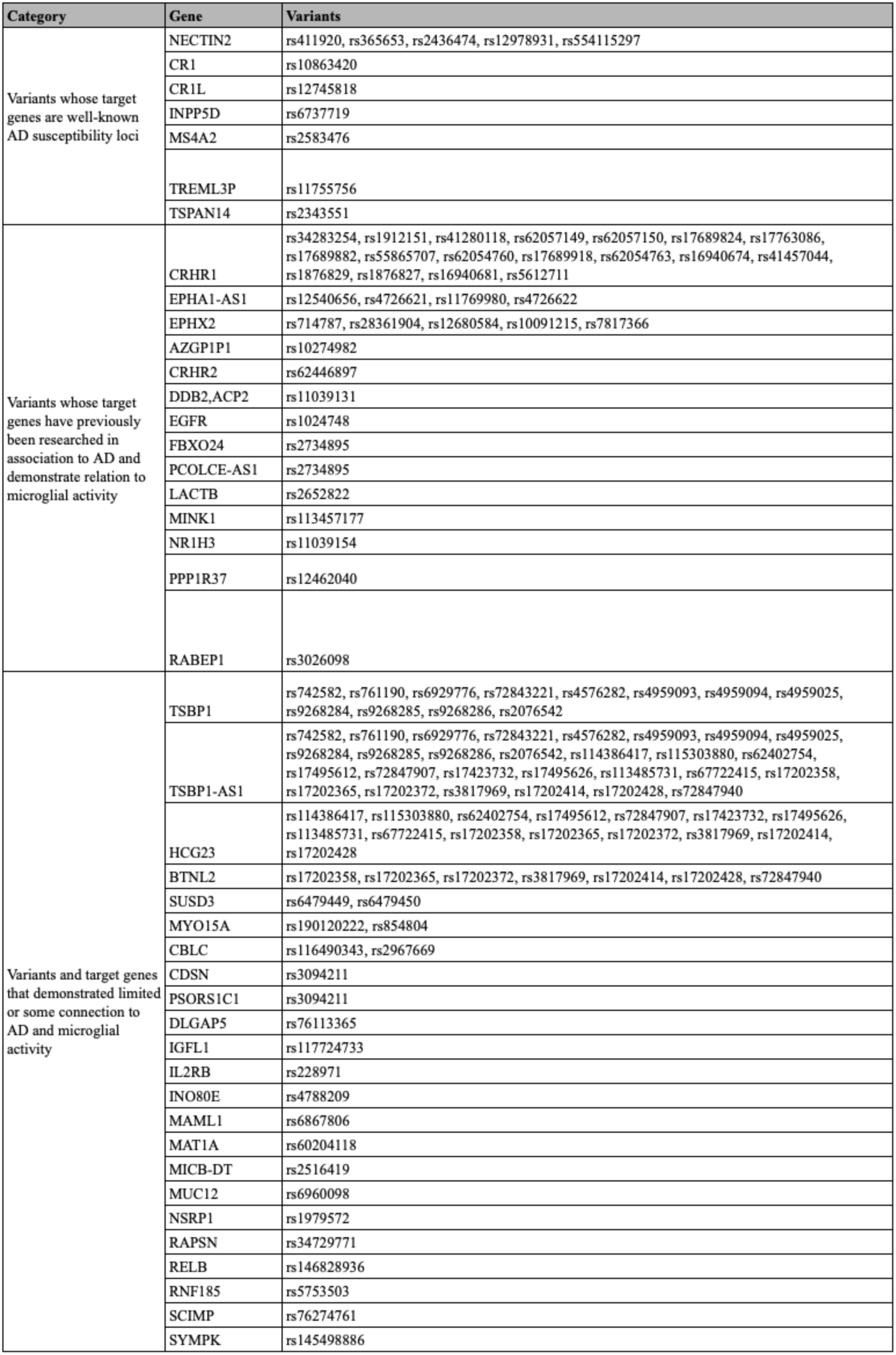
Complete List of 155 Top-Performing Variants and Their Target Genes. Genes are divided into three categories based their association to AD and microglia: 1) variants whose target genes are well-known AD susceptibility loci, 2) variants whose target genes have previously been researched in association to AD and demonstrate relation to microglial activity, and 3) variants and target genes that demonstrated limited or some connection to AD and microglial activity.

