## Supplementary figure and table for "S-BEAM: A Semi-Supervised Ensemble Approach to Rank Potential Causal Variants and Their Target Genes in Microglia for Alzheimer’s Disease"

### Supplementary Materials

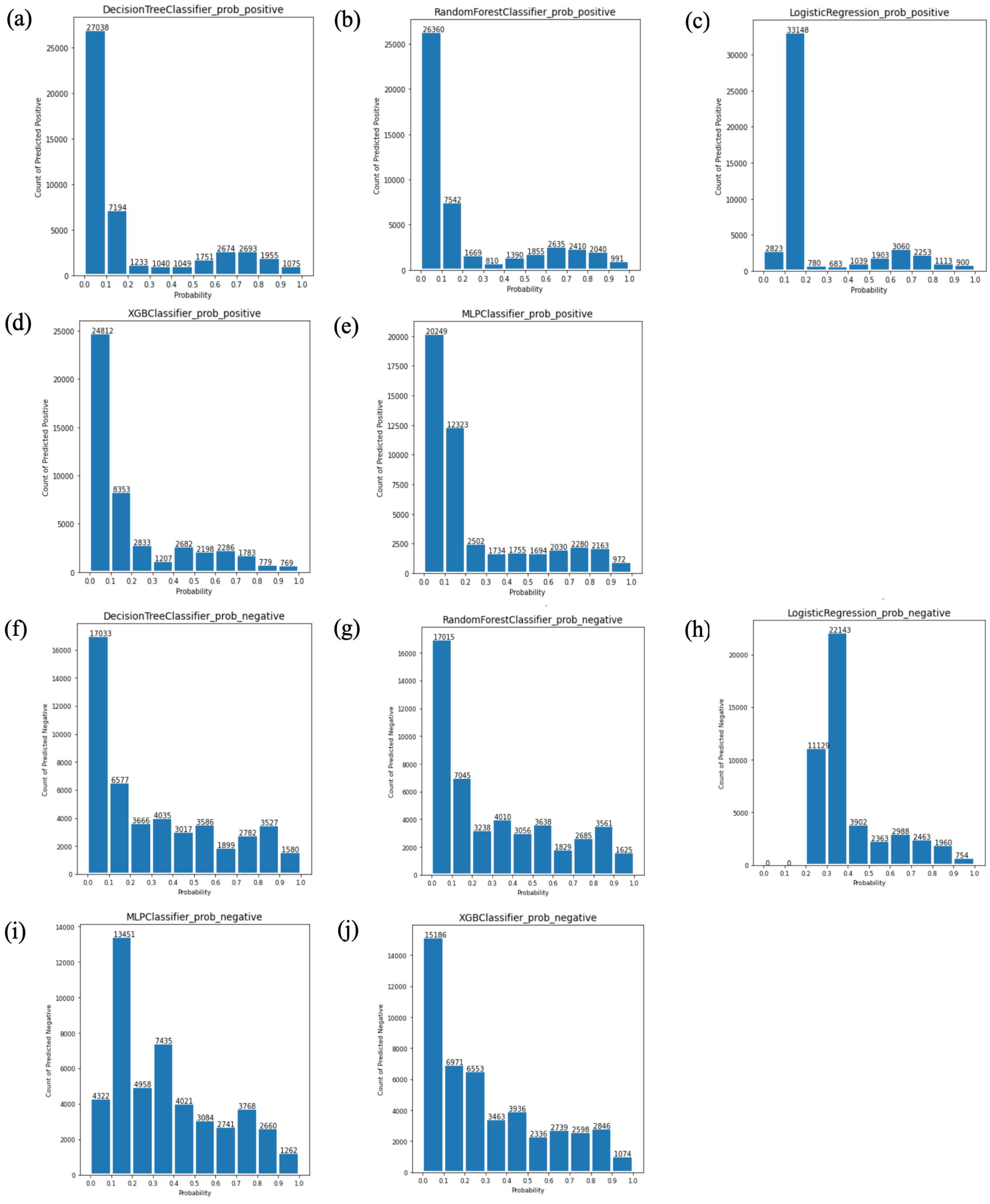


**Supplementary Figure 1: Bin Charts for Individual PU and NU Classifiers.** These histograms show the distribution of predicted posterior probabilities across 10 bins for each of the 5 base classifiers implemented in our model.

| Category | Gene | Variants |
|---|---|---|
| Variants whose target genes are well-known AD susceptibility loci | NECTIN2 | rs411920, rs365653, rs2436474, rs12978931, rs554115297 |
| | CR1 | rs10863420 |
| | CR1L | rs12745818 |
| | INPP5D | rs6737719 |
| | MS4A2 | rs2583476 |
| | TREML3P | rs11755756 |
| | TSPAN14 | rs2343551 |
| Variants whose target genes have previously been researched in association to AD and demonstrate relation to microglial activity | CRHR1 | rs34283254, rs1912151, rs41280118, rs62057149, rs62057150, rs17689824, rs17763086, rs17689882, rs55865707, rs62054760, rs17689918, rs62054763, rs16940674, rs41457044, rs1876829, rs1876827, rs16940681, rs5612711 |
| | EPHA1-AS1 | rs12540656, rs4726621, rs11769980, rs4726622 |
| | EPHX2 | rs714787, rs28361904, rs12680584, rs10091215, rs7817366 |
| | AZGP1P1 | rs10274982 |
| | CRHR2 | rs62446897 |
| | DDB2,ACP2 | rs11039131 |
| | EGFR | rs1024748 |
| | FBXO24 | rs2734895 |
| | PCOLCE-AS1 | rs2734895 |
| | LACTB | rs2652822 |
| | MINK1 | rs113457177 |
| | NR1H3 | rs11039154 |
| | PPP1R37 | rs12462040 |
| | RABEP1 | rs3026098 |
| Variants and target genes that demonstrated limited or some connection to AD and microglial activity | TSBP1 | rs742582, rs761190, rs6929776, rs72843221, rs4576282, rs4959093, rs4959094, rs4959025, rs9268284, rs9268285, rs9268286, rs2076542 |
| | TSBP1-AS1 | rs742582, rs761190, rs6929776, rs72843221, rs4576282, rs4959093, rs4959094, rs4959025, rs9268284, rs9268285, rs9268286, rs2076542, rs114386417, rs115303880, rs62402754, rs17495612, rs72847907, rs17423732, rs17495626, rs113485731, rs67722415, rs17202358, rs17202365, rs17202372, rs3817969, rs17202414, rs17202428, rs72847940 |
| | HCG23 | rs114386417, rs115303880, rs62402754, rs17495612, rs72847907, rs17423732, rs17495626, rs113485731, rs67722415, rs17202358, rs17202365, rs17202372, rs3817969, rs17202414, rs17202428 |
| | BTNL2 | rs17202358, rs17202365, rs17202372, rs3817969, rs17202414, rs17202428, rs72847940 |
| | SUSD3 | rs6479449, rs6479450 |
| | MYO15A | rs190120222, rs854804 |
| | CBLC | rs116490343, rs2967669 |
| | CDSN | rs3094211 |
| | PSORS1C1 | rs3094211 |
| | DLGAP5 | rs76113365 |
| | IGFL1 | rs117724733 |
| | IL2RB | rs228971 |
| | INO80E | rs4788209 |
| | MAML1 | rs6867806 |
| | MAT1A | rs60204118 |
| | MICB-DT | rs2516419 |
| | MUC12 | rs6960098 |
| | NSRP1 | rs1979572 |
| | RAPSN | rs34729771 |
| | RELB | rs146828936 |
| | RNF185 | rs5753503 |
| | SCIMP | rs76274761 |
| | SYMPK | rs145498886 |

**Supplementary Table 1: Complete List of 155 Top-Performing Variants and Their Target Genes.** Genes are divided into three categories based their association to AD and microglia: 1) variants whose target genes are well-known AD susceptibility loci, 2) variants whose target genes have previously been researched in association to AD and demonstrate relation to microglial activity, and 3) variants and target genes that demonstrated limited or some connection to AD and microglial activity.
